# The Neuroanatomy of the Hawaiian Bobtail Squid Juvenile Bacterial Light Organ

**DOI:** 10.64898/2026.05.15.725553

**Authors:** Alice Breaux Walker, Elizabeth Victoria Xiu Widun, Elizabeth Anne Chapman Heath-Heckman

## Abstract

**Background:** Recent studies have shown that symbiotic bacteria can have drastic effects on host neurobiology, but few simple, accessible models currently exist in which to study these interactions. Hawaiian bobtail squid (*Euprymna scolopes*) participate in a binary symbiosis with the bacterium *Vibrio fischeri*, a population of which resides in a specialized hindgut-derived organ called the light organ. Upon colonization by *V. fischeri*, the light organ undergoes transcriptional changes that suggest neurons are among the cell types impacted by the initiation of symbiosis, but the nascent light organ’s innervation has remained uncharacterized.

**Results:** The light organ-associated nervous system (LONS) in hatchling *E. scolopes* is a remarkably complex segment of the peripheral nervous system. The LONS is largely plexiform and originates from two primary nerves connected by a local commissure. The abundance of synapsin-like immunoreactivity indicates that the lobe plexus is highly interconnected. We also highlight a small number of serotonin-like immunoreactive neurites innervating the anterior appendages that are poised to be directly impacted by symbiont-driven post-embryonic development. Finally, we present evidence that a limited but morphologically diverse population of neurons reside within the light organ and are often located near internal symbiont-interacting structures.

**Conclusion:** Our results show that the LONS is structurally and molecularly complex and exhibits traits characteristic of an immature nervous system, suggesting that it may undergo substantial post-embryonic refinement. This initial characterization of the LONS provides a foundation from which to investigate how beneficial bacterial symbionts affect host peripheral neurobiology in a tractable model system.

## 1 Background

The highly advanced nervous systems of coleoid cephalopods provide a unique opportunity to understand the common principles that govern nervous system organization and development [1, 2]. Though the majority of research efforts have focused on the cephalopod central nervous system and the independent evolutionary origin of their large, centralized brains [3], the cephalopod peripheral nervous system (PNS) remains understudied. Perhaps the most thoroughly characterized segment of the cephalopod PNS is the neuroanatomy of the octopus arm. Each arm contains a large axial nerve cord akin to the vertebrate spinal cord as well as four intramuscular nerve cords that carry sensory and motor information [4]. Further, each of hundreds of suckers has a dedicated peripheral ganglion [5] and chemotactile receptors that line the sucker sensory epithelium [6]. Together, these features allow the octopus arm to locally integrate sensory information and perform a vast repertoire of autonomous behaviors [4]. Less studied, however, is the segment of the cephalopod PNS that innervates the visceral organs, including the ink sac, gut, and related structures. The innervation of octopus viscera was initially described as being composed of numerous small fibers in plexiform arrangements originating from two visceral nerves [7]. It has been shown more recently that the gastric ganglion, the largest cephalopod visceral ganglion, is both structurally and neurochemically complex in *Octopus vulgaris* (the common octopus), consisting of a dense neuropil surrounded by neuron cell bodies expressing a wide array of neuroactive substances [8]. Similar attention has not yet been given to visceral PNS structures in other cephalopods.

The enteric nervous system (ENS) is the division of the PNS that innervates the gastrointestinal tract and governs gut motility, blood flow, and the secretion of digestive and immune factors, all of which contribute to a luminal environment that supports the gut microbiome [9]. The ENS and the gut microbiome have wide-ranging, often bidirectional impacts on one another. Mouse enterochromaffin cells and enteric neurons release serotonin (5-hydroxytryptamine, 5-HT) into the intestinal lumen, which reduces the expression of virulence genes in the bacterial pathogen *Citrobacter rodentium* [10]. Hydra have been shown to influence their microbiome population structure by using the neuron-secreted peptide NDA-1 to inhibit bacterial growth [11]. In zebrafish, disruption of genes associated with the development of enteric neurons and glia (*sox10*, *ret*) results in not only gut motility defects, but also altered gut microbiome composition [12, 13]. Conversely, the gut microbiome often mediates normal ENS development and maintenance. Beneficial gut bacteria stimulate the production of 5-HT in the ENS, resulting in the maturation of enteric neurons via 5-HT_4_R activation [14]. The absence of a complex gut microbiome in germ-free mice, however, yields defects in microglial immune functionality, decreased gut motility, and reduced neuronal density the myenteric plexus [15, 16]. The precise mechanisms by which non-pathogenic bacteria exert their effects on host peripheral neurobiology, however, remain poorly understood. Current models of ENS-microbe interactions typically require host animals to be raised in a germ-free environment due to the gut’s permissiveness to colonization by a wide array of microorganisms, highlighting the need for a simpler, easily manipulable model. Thus, the squid-*Vibrio* system represents a potent model system to interrogate beneficial microbe-nervous system interactions by virtue of its specificity.

The Hawaiian bobtail squid (*Euprymna scolopes*) participates in a binary mutualism with the bioluminescent marine bacterium *Vibrio fischeri* in which *E. scolopes* provides *V. fischeri* with nutrients and shelter in exchange for the production of light [17, 18]. The squid harnesses bacterial bioluminescence to camouflage itself from predators via a process called counterillumination [19]. *E. scolopes* houses its symbionts in a novel organ called the light organ, which develops from the hindgut-ink sac complex and is located immediately dorsal to the hindgut [20]. Since *E. scolopes* obtain their symbionts after hatching, the light organ is highly specialized to not only shelter *V. fischeri*, but also acquire it from the water column. To facilitate the process of symbiont acquisition, the hatchling light organ possesses two pairs of appendages that project from the superficial ciliated epithelium [17] (Figures 1b-c). A subset of the cilia beat in a metachronal rhythm to entrain environmental bacteria toward the surface of the light organ, where *V. fischeri* form aggregates in mucus secreted by the host [21, 22]. *V. fischeri* then begins its approximately 150 μm migration through one of six pores, ciliated ducts, and antechambers, followed by a physical bottleneck before reaching the deep crypts (Figure 1), where the symbiont will ultimately reside, reproduce, and luminesce [23]. We herein refer to these light organ structures (ducts, antechambers, and deep crypts) collectively as the internal symbiont-interacting structures (ISS).

**FIGURE 1.**
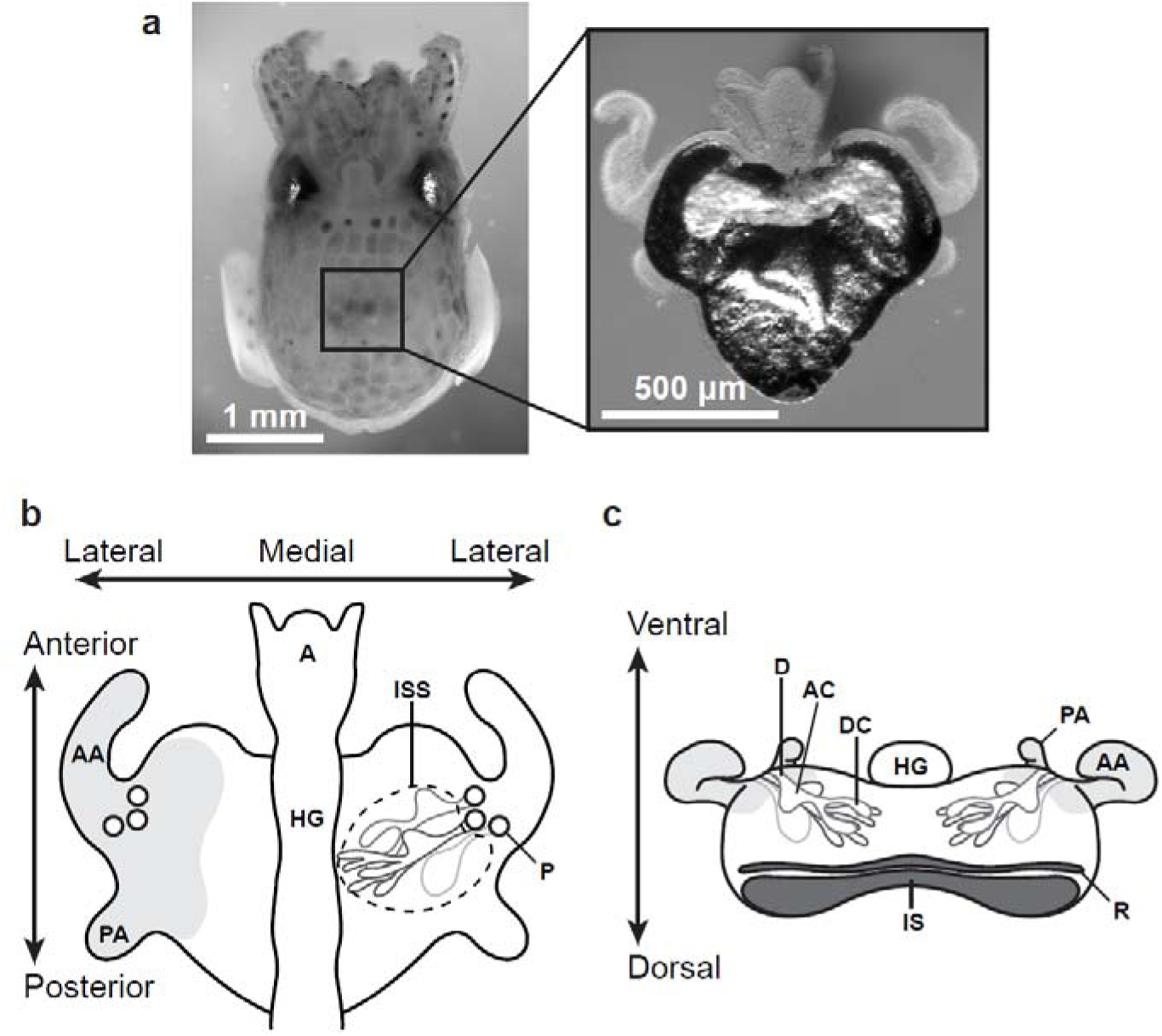
Overview of light organ anatomy. (**a**) A hatchling Hawaiian bobtail squid, *Euprymna scolopes* (left), and its light organ (right) pictured from their ventral aspects. (**b-c**) Diagrams of relevant light organ internal structures and functional orientation axes. Regions shaded in light grey represent the superficial ciliated epithelium (**b**) Ventral aspect diagram of the light organ. The left half depicts surface features of the light organ, while the right half is cut away to reveal three sets of ducts (D), antechambers (AC), and deep crypts, collectively referred to as internal symbiont-interacting structures (ISS). The hindgut (HG) contacts the light organ ventrally along its midline. (**c**) Anterior aspect diagram of the light organ. The main body of the light organ is cut away to reveal the positions of the D, AC, and DC relative to the reflector (R) and ink sac (IS). A, anus; AA, anterior appendage; AC, antechamber; D, duct; DC, deep crypt; HG, hindgut; IS, ink sac; ISS, internal symbiont-interacting structures; P, pore; PA, posterior appendage; R, reflector.

Upon colonizing the deep crypts, *V. fischeri* signals to the host via microbe-associated molecular patterns (MAMPs) which causes the ciliated epithelium to undergo apoptosis, resulting in the complete regression of the appendages after approximately 4 days [24–26]. Additionally, initial detection of *V. fischeri* at the superficial ciliated epithelium results in transcriptional changes throughout the hatchling light organ that serve to facilitate efficient symbiont colonization, including the regulation of genes associated with visual perception, phototransduction, anion transmembrane transport, and chloride transport [27, 28]. These data suggest that *E. scolopes* neurobiology is affected by the onset of its symbiosis with *V. fischeri*, but little is known about the innervation of the light organ. This study serves to establish a baseline understanding of the branch of *E. scolopes* PNS that innervates the light organ, which we have termed the light organ-associated nervous system (LONS), in hatchling animals that are not yet exposed to *V. fischeri*.

## 2 Results

### 2.1 Light organ-associated nervous system macrostructures are revealed by serotonin immunoreactivity

Our first objective was to visualize the major components of the LONS by using immunocytochemistry. Due to the remarkably high abundance of acetylated alpha tubulin in the superficial ciliated epithelium, labeling this common marker of neuronal processes proved ineffective for visualizing the LONS. Instead, we targeted neuroactive substances that are abundant and show robust staining in the PNS of other cephalopods. We selected a commercially available polyclonal antibody shown to specifically bind 5-HT in the nervous systems of *Idiosepius notoides* [29] (southern pygmy squid), *O. vulgaris* [30], and nudibranch mollusks [31]. 5-HT-like immunoreactivity (-lir) revealed an elaborate meshwork of innervation throughout the light organ which could be divided into the three main structures that we have defined herein (Figure 2). The ventromedial nerves (VMNs; Figure 2a) are a pair of large, bilaterally symmetric nerves that appear to enter the light organ from its anterodorsal aspect (Figure 2e, asterisk) and travel ventrally. Upon reaching the ventral surface of the lobe, each VMN then projects posteriorly (Figures 2b-e). The VMNs are connected near the anterior margin of the light organ by the ventral commissure (VC; Figures 2a and c), a similarly sized neurite bundle that spans the midline of the light organ. Each VMN runs along the anterior-posterior axis roughly parallel to the hindgut and branches into smaller neurite bundles that make up the lobe plexus (LP; Figure 2a). Neurite bundles that compose the LP initially travel dorsally from the VMN and spread across the light organ lobe near its boundary with the ink sac (Figures 2d-e). LP neurite bundles further defasciculate as they project ventrally to form a complex meshwork of innervation throughout each light organ lobe. The neurite bundles exhibit an apparently disordered, plexiform arrangement, but they appear to decrease in diameter from the dorsal LP to the ventral LP (Figures 2b-e). Each lobe also contains a central region with an absence of LP innervation, which corresponds to the space occupied by the ISS (Figures 2c-d). Notably, we were unable to identify any 5-HT-lir neuron cell bodies in the light organ, but it is possible that they were obscured by the exuberance of 5-HT-lir neurites in the LONS. Taken together, the gross organization of the LONS suggests that the light organ lobes and the structures necessary to initiate and establish the squid-*Vibrio* partnership may be under the control of a robust segment of the host’s PNS.

**FIGURE 2.**
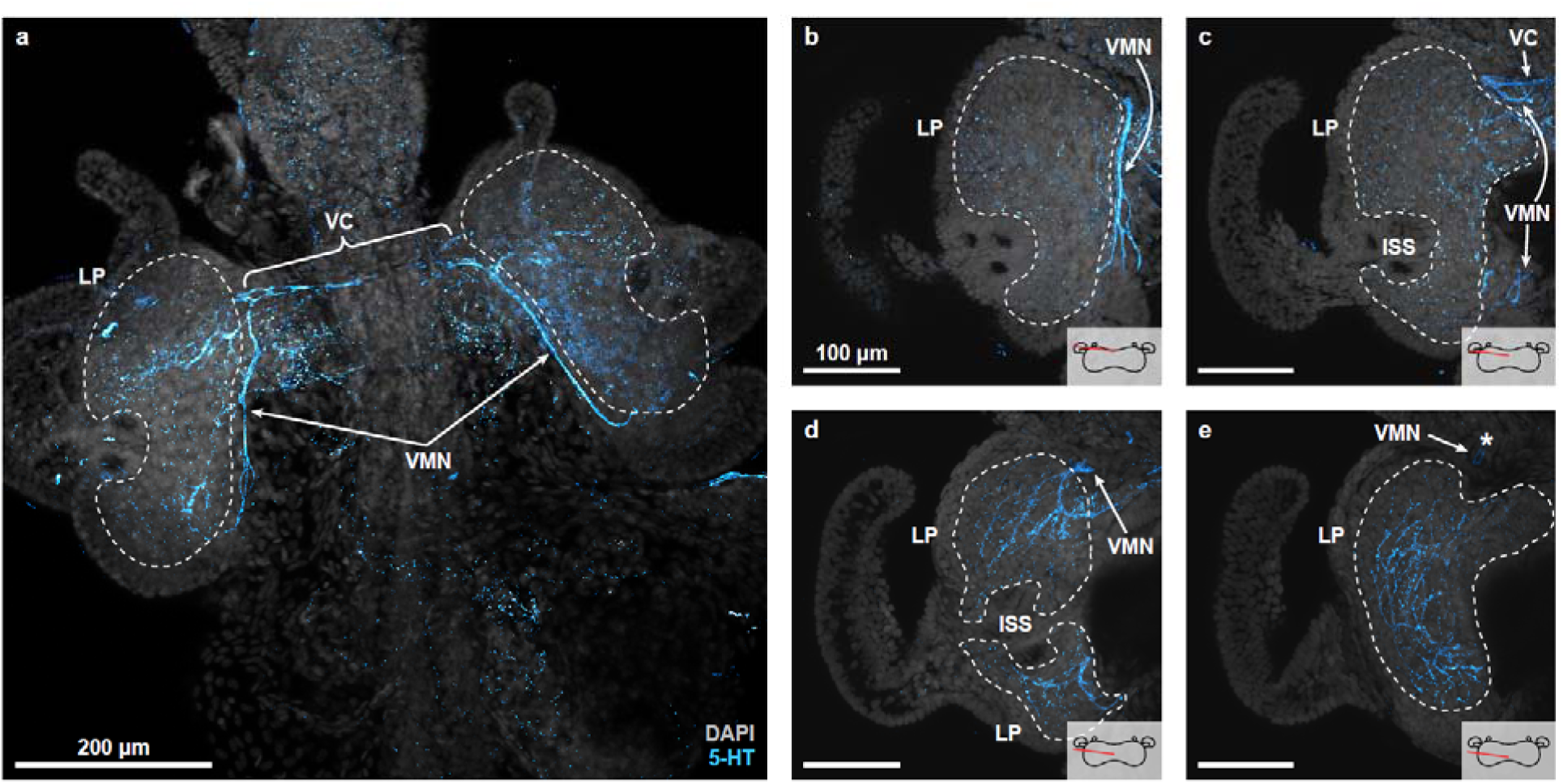
Macrostructures of the light organ-associated nervous system are revealed by serotonin immunostaining (5-HT, cyan). (**a**) Maximum intensity projection showing the three major segments of the light organ-associated nervous system (LONS). Two ventromedial nerves (VMNs) are joined by a ventral commissure (VC), which spans the midline of the light organ. Each VMN gives rise to a lobe plexus (LP) that spreads laterally throughout the remainder of the light organ. (**b-e**) One half of the LONS depicted in segments along the dorsal-ventral (D-V) axis by dividing a confocal Z-stack into four equally spaced maximum intensity projections. The approximate D-V position of each projection is indicated by a red line in the inset schematic. (**b**) Most ventrally, the VMN extends along the anterior-posterior axis, ramifying into multiple neurite bundles. Individual neurites comprising the ventral LP are also visible. (**c**) The VMN connects to the VC at its anterior end and projects dorsally at the anterior and posterior margins of the light organ. Small neurite bundles are present in the medial LP, while individual neurites innervate the region surrounding the internal symbiont-interacting structures (ISS). (**d**) At this depth, the LP is composed of larger neurite bundles that defasciculate from the VMN. LP projections surround the ISS. (**e**) Larger neurite bundles are visible in the dorsal LP, where the LONS borders the reflector. A branch of the VMN appears to terminate at the anterior margin of the light organ (asterisk). Scale bars are consistent within (**b-e**). Nuclei are stained with DAPI (grey). 5-HT, serotonin; ISS, internal symbiont-interacting structures; LP, lobe plexus; VC, ventral commissure; VMN, ventromedial nerve.

### 2.2 The lobe plexus contains NeuN-lir cell bodies

To visualize the spatial distribution of putative neuron cell bodies in the LONS, we stained light organs with a commercially available polyclonal antibody raised against the neuron-specific nuclear antigen NeuN for the first time in *E. scolopes*. NeuN (also known as Rbfox-3) is an RNA-binding protein that regulates alternative pre-mRNA splicing in post-mitotic neurons and is therefore a widely used as a marker for neuronal nuclei [32–34]. The polyclonal antibody we selected effectively labels neuron cell bodies in the PNS of *O. vulgaris* [8]. We identified NeuN-lir cells by a characteristic perinuclear staining pattern [35–37] (Figure 3a). The precise location of NeuN-lir cells varied between light organs, so we aggregated observations from 10 individuals and performed two-dimensional kernel density estimates (2D-KDEs) to visualize probability densities in the horizontal and transverse planes of the light organ using a normalized coordinate system [38] (Figures 3c-e). “Normalized units” are used to refer to distances within this coordinate system. The distribution of NeuN-lir cells in each light organ used to generate the 2D-KDEs can be found in Supplemental Figure S1 (Additional file 1).

**FIGURE 3.**
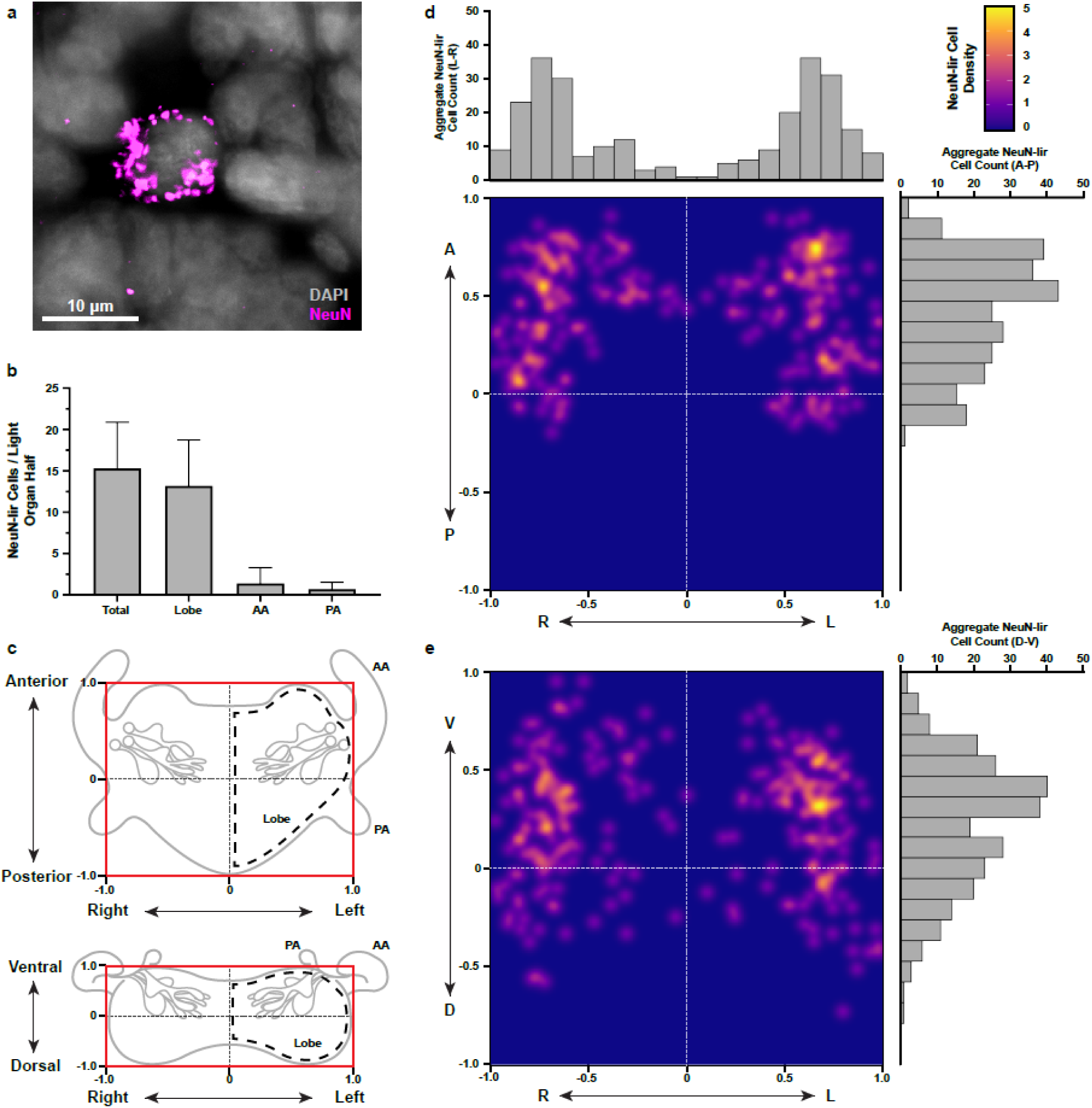
Spatial distribution of NeuN-like immunoreactive cells. (**a**) Immunocytochemistry reveals light organ cells with perinuclear NeuN-like immunoreactivity (NeuN-lir, magenta). Nuclei are stained with DAPI (grey). (**b**) NeuN-lir cells are more abundant in the light organ lobes than the anterior appendages (AA) and posterior appendages (PA). Error bars represent standard error of the mean; *n* = 10 biological replicates. (**c**) Normalized coordinate planes for ventral (top) and anterior (bottom) views of the light organ. Red boxes bounding each light organ diagram correspond to the extreme values of the coordinate planes depicted in (**d-e**). (**d-e**) Two-dimensional kernel density estimates of NeuN-lir cell positions in the frontal (**d**) and transverse (**e**) planes. Accompanying histograms show the total number of NeuN-lir cell observations distributed across each axis for *n* = 10 biological replicates. NeuN-lir cells are most abundant in regions corresponding to the lobe plexuses. Symmetric high-density regions in each lobe spatially overlap with the position of internal symbiont-interacting structures. AA, anterior appendage; A, anterior; D, dorsal; L, left; P, posterior; PA, posterior appendage, R, right; V, ventral.

NeuN-lir cells were also observed in the appendages but were excluded from the 2D-KDEs due to the variable positioning of the appendages during imaging. Instead, we counted the total number of NeuN-lir cells in one half of each light organ for statistical independence and grouped them according to whether they occurred in the lobe, the anterior appendage (AA), or the posterior appendage (PA), which showed that the majority of putative neuron cell bodies in the LONS reside in the lobes (Figure 3b).

NeuN-lir cells in the main body of the light organ were most often located near the anterior periphery of the LP, but very few NeuN-lir cells were observed at the extreme margins of the lobes (Figure 3d). The lower frequency of NeuN-lir cell observations in the posterior and dorsal portions of the LONS can be attributed to the location of the reflector and ink sac (Figures 1c and 3d-e). Segmenting the light organ along the dorsoventral axis revealed that NeuN-lir cells were more likely to be observed in the ventral half of the LP (Figure 3e). Areas of high NeuN-lir cell density (i.e., probability density ≥ 3.0) were located within 0.2 normalized units of the ISS periphery, and high-density areas frequently overlapped with the ISS in the frontal plane (Figures 3c-e).

The LONS contains relatively few NeuN-lir cell bodies in comparison to its abundance of neurites (Figure 2a). We also did not observe distinct clusters of NeuN-lir cell bodies within a single light organ that would indicate the presence of peripheral ganglia in the LONS (Additional file 1: Figure S1). From this, we conclude that the light organ is at least partially extrinsically innervated, though the origin of this innervation has yet to be determined.

### 2.3 Golgi-Cox staining reveals cells consistent with multiple neuron morphologies

In the preceding experiments, we observed neuronal processes and putative neuron cell bodies in isolation from one another. Therefore, to visualize entire LONS neurons and characterize their morphologies, we developed a novel Golgi-Cox staining protocol by combining multiple established techniques [39–41] to optimally stain LONS cells in *E. scolopes* hatchlings. Golgi-Cox staining is an inexpensive technique that utilizes heavy metal precipitates to impregnate a small subset of cells to reveal intricate neuronal arbors with exquisite clarity, making it well-suited for studying the morphology of individual neurons within the highly elaborate LP.

We observed neurons with three main morphological profiles across 15 successfully stained (Golgi-Cox^+^) light organs: multipolar cells (Figures 4a and b), bipolar cells (Figure 4c), and unipolar cells (Figure 4d). Golgi-Cox staining revealed a prominent subset of bipolar neurons, with two primary neurites projecting from opposing poles of the cell body [42] (Figures 4c-d). The length and branching structure of these neurites varies considerably within individual light organs. When viewing the light organ from its ventral aspect, the cell bodies of bipolar neurons with long primary neurites were located within a 20 μm periphery of the ISS, with some cells overlapping with the ISS in the frontal plane. Primary projections from long bipolar cells appeared to encircle the ISS (Figure 4c). The remaining bipolar neurons have relatively shorter primary neurites that do not exhibit substantial branching distal to the neuron cell body. These short bipolar cells are distributed throughout the LP and do not exhibit a distinct pattern of localization. (Figures 3d-e).

**FIGURE 4.**
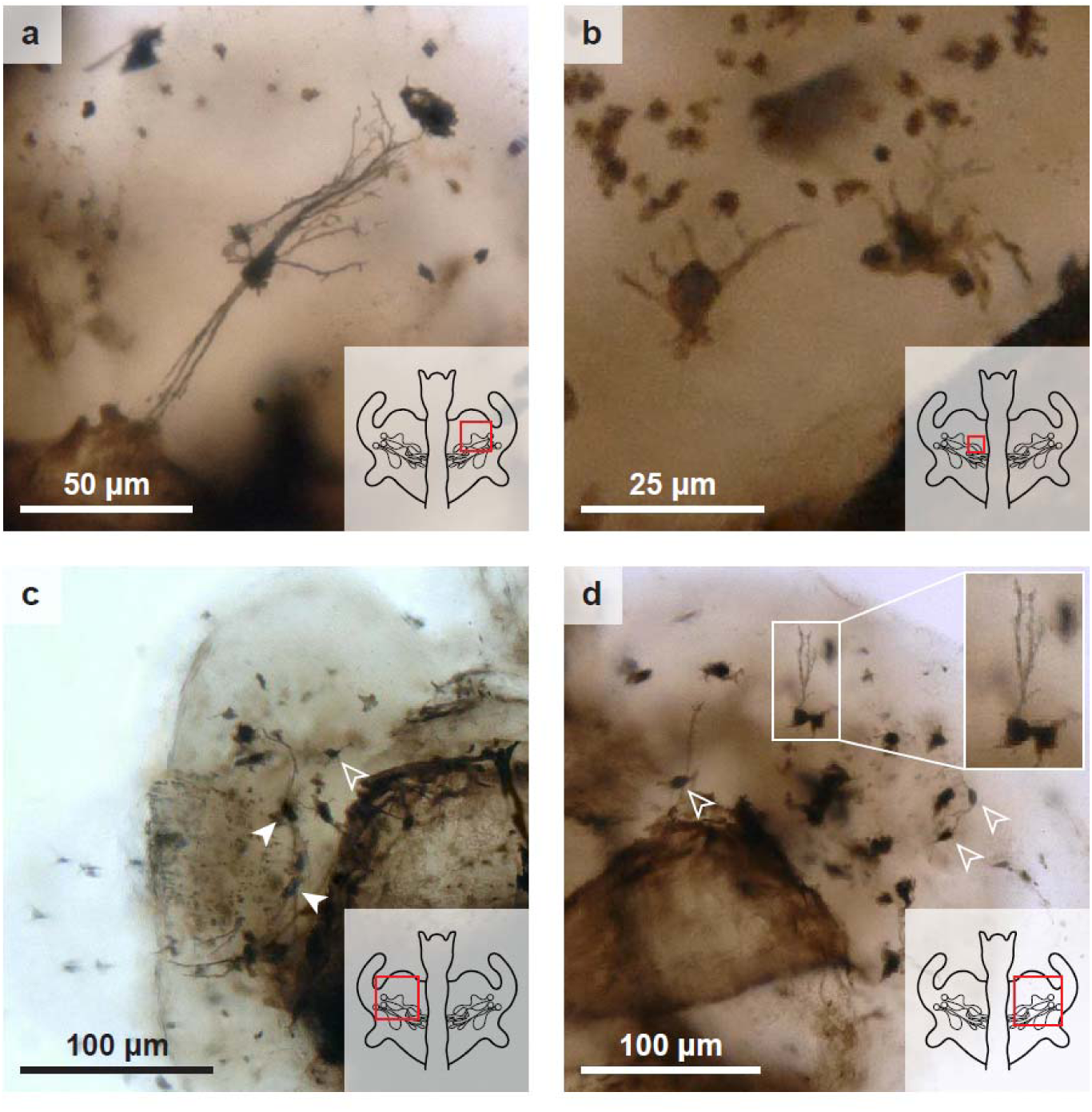
Golgi-Cox staining reveals cells with neuronal morphology in the lobe plexus. (**a**) A large multipolar neuron with double-bouquet-like morphology. (**b**) Two short multipolar neurons with short projections surrounding the internal symbiont-interacting structures (ISS). (**c-d**) Long bipolar neurons (closed arrowheads), short bipolar neurons (open arrowheads), and a unipolar neuron (inset) in the lobe plexus (LP). (**c**) Bipolar neurons in the LP with projections that form a basket-like pattern surrounding the ISS. (**d**) Short bipolar neurons and a unipolar neuron stationed near ISS. The region of the light organ depicted in each panel is indicated by a red square in the inset schematic.

Multipolar neurons in the LONS exhibit a similar degree of morphological variation to the bipolar cells (Figures 4a-b). The majority of multipolar neurons have diminutive neurite arbors in comparison to other Golgi-Cox^+^ cells, and they appear evenly distributed among all stained cells. These neurons frequently have irregular, compact neurite arbors that do not extend a significant distance away from the cell body (Figure 4b). We did, however, observe a select few multipolar neurons that displayed an arrangement of neurites that resembles the double bouquet neurons of the primate cerebral cortex [43] (Figure 4a). Double bouquet-like LONS neurons have more than two primary neurites that project from the cell body, but the neurites all originate from opposing poles of the cell body, forming two discrete tufts of neurites. Only three Golgi-Cox^+^ cells across 15 light organs displayed double bouquet-like morphology, indicating that this may be a low-frequency cell type in the LONS. These cells were consistently located in the lateral anterior quadrant of the light organ lobe. Notably, we never observed more than one double bouquet-like neuron in a single light organ lobe, suggesting that this rare cell type may have a specialized function in the LONS.

Unipolar neurons were the least prominent of the three morphological categories, with only a single representative cell across 15 light organs (Figure 4d, asterisk). This cell has only one visible primary neurite which ramifies into three prominent branches, which is reminiscent of the primary branching pattern of some unipolar brush cells (UBCs) in the primate cerebellum [44, 45]. However, this unipolar cell appears to lack terminal dendriole brushes, making it structurally distinct from the primate UBC. The morphological heterogeneity of LONS neurons suggests that they may also exhibit significant genetic diversity that could be leveraged to target subpopulations of these cells in future studies.

### 2.4 The LONS is neurochemically heterogeneous

To further interrogate the neurochemical makeup of the LONS, we employed a commercially available polyclonal antibody raised against FMRFamide, a common invertebrate neuropeptide first discovered in the venus clam *Macrocallista nimbosa* [46] and later identified in other mollusks, including cephalopods [47]. This polyclonal antibody exhibits cross-reactivity with other FMRFamide-related peptides (FaRPs) containing either a YLRFamide or FxRFamide C-terminus [48, 49], so we herein refer to immunoreactivity attributed to this antibody as FaRP-like. Immunocytochemistry revealed widespread FaRP-lir throughout the LONS (Figure 5c). FaRP-lir in the LONS spatially overlaps with 5-HT-lir (Figure 5a), but each neuroactive substance has a distinct staining pattern (Figures 5a and c). The majority of FaRP-lir neurites in the LP were observed in the posterior half of the light organ and closer to the midline. 5-HT-lir neurites, however, were distributed more evenly along dorsal-ventral axis and projected farther toward the periphery of the light organ lobe.

**FIGURE 5.**
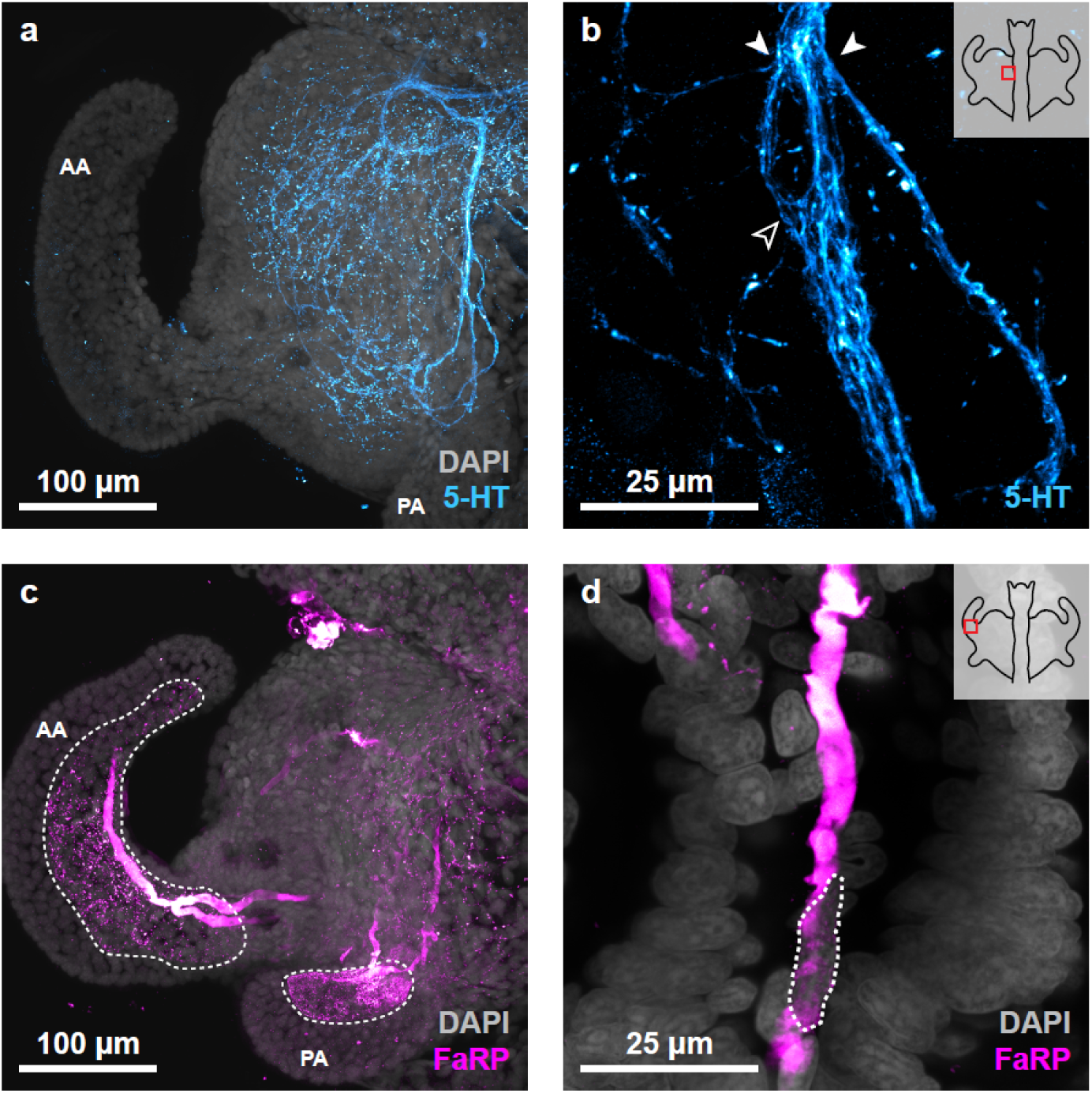
Neuroactive substances localize to different light organ structures. (**a**) Maximum intensity projection of serotonin (5-HT, cyan) immunostaining in one half of the light organ-associated nervous system (LONS) reveals the ventromedial nerve (VMN), ventral commissure, and lobe plexus (LP). (**b**) 5-HT-like immunoreactive (5-HT-lir) neurites in the VMN. Small neurite bundles and single neurites defasciculate from (closed arrowheads) and rejoin (open arrowhead) the VMN. (**c**) Maximum intensity projection of FMRFamide-related peptide (FaRP, magenta) immunostaining in one half of the LONS reveals neurites in the medial LP, as well as blood vessels that invade the light organ lobes and appendage blood sinuses (dashed outline). (**d**) FaRP-lir in a prominent blood vessel near the base of the anterior appendage (AA). The nucleus of a hemocyte being trafficked through the blood vessel is also visible (dashed outline). The regions of the light organ depicted in (**b**) and (**d**) are indicated by a red square in the inset schematic. Nuclei are stained with DAPI (grey). 5-HT, serotonin; AA, anterior appendage; FaRP, FMRFamide-related peptide; PA, posterior appendage.

Additionally, 5-HT-lir and FaRP-lir highlight distinct features of the LONS (Figures 5b and d). 5-HT-lir strongly labeled the major neurite bundles of the LONS, the VC and VMNs (Figure 5b). This revealed locations along the VMN where small bundles of neurites defasciculated (closed arrowheads) and rejoined (open arrowheads) the neurite bundle. Only sparse FaRP-lir was observed in locations that correspond to large neurite bundles (Figure 5c). This pattern was also observed in the LP, with 5-HT-lir revealing both individual neurites and the large neurite bundles of the dorsal LP (Figure 2), while FaRP-lir was limited to labeling small bundles throughout the LP (Figure 5c). Interestingly, FaRP-lir also revealed non-neuronal structures in the light organ. Diffuse FaRP-lir was consistently observed in both the vasculature that terminates in the anterior and posterior appendages, as well as the blood sinuses of the appendages (Figures 5c-d).

### 2.5 The lobe plexus is highly interconnected

Having identified neuronal processes that appear to terminate within the LONS, we next aimed to identify which cells were the targets of this innervation. To do so, we used a commercially available synapsin-1 (Syn) monoclonal antibody to visualize the location and distribution of synapses in the LONS. Syn-lir was punctate and widespread throughout the LP, overlapping with 5-HT-lir and FaRP-lir regions (Figures 6a and e). Syn-lir was less frequently observed near the margins of the LP, corresponding to the decreased abundance of 5-HT-lir and FaRP-lir in this portion of the light organ (Figures 5a and c). Additionally, co-labeling FaRPs and Syn revealed that the anterior half of the LP contains substantial Syn-lir but a paucity of FaRP-lir (Figure 6e). This same region, however, showed significant overlap between 5-HT-lir and Syn-lir (Figure 6a), supporting the claim that the LP may contain at least two distinct populations of neurons that are distinguishable by the neuroactive substances they contain.

**FIGURE 6.**
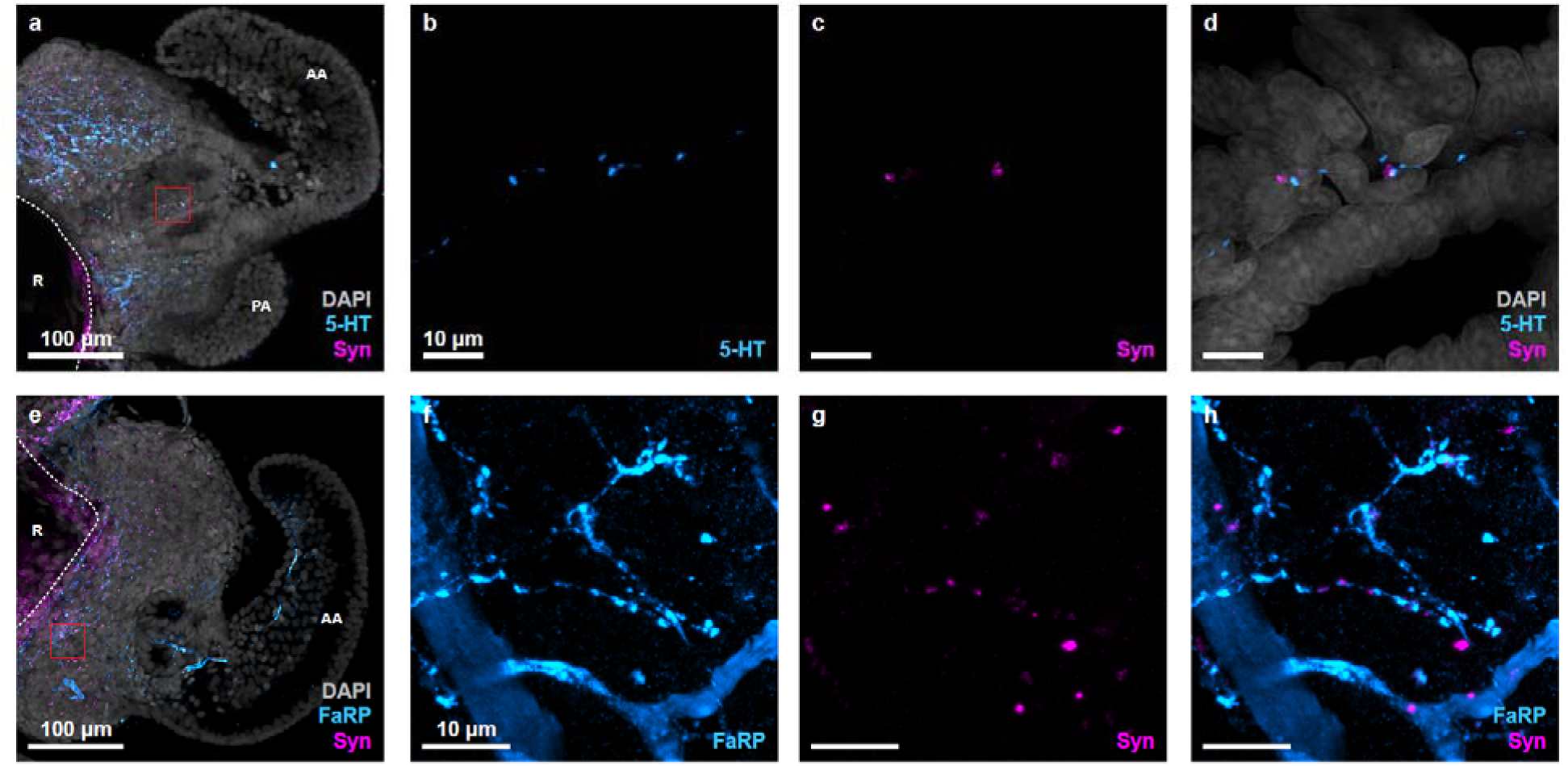
The lobe plexus is highly interconnected. (**a-d**) The light organ-associated nervous system immunostained for serotonin (5-HT, cyan) and synapsin-1 (Syn, magenta). (**a**) Syn-like immunoreactive (Syn-lir) regions of the lobe plexus (LP) substantially overlap with 5-HT-lir regions. (**b-d**) 5-HT-lir (**b**) and Syn-lir (**c**) localize to a single neurite that travels between two ducts. (**d**) Merge of (**b**) and (**c**). (**e-h**) The LONS immunostained for FMRFamide-related peptides (FaRP, cyan) and Syn (magenta). (**e**) The Syn-lir region of the LP only partially overlaps with FaRP-lir regions. Most overlap occurs in the medial LP. (**f-h**) FaRP-lir (**f**) and Syn-lir (**g**) localize to single neurites and light organ vasculature in the medial LP. (**h**) Merge of (**f**) and (**g**). Panels (**b-d**) and (**f-h**) were captured at higher magnification than (**a**) and (**e**), and the regions depicted are indicated by a red square in (**a**) and (**e**) respectively. Scale bars are consistent within (**b-d**) and (**f-h**). Nuclei are stained with DAPI (grey). 5-HT, serotonin; AA, anterior appendage; FaRP, FMRFamide-related peptide; PA, posterior appendage; R, reflector; Syn, synapsin-1.

At smaller scales, 5-HT and FaRPs had different spatial relationships with Syn (Figures 6b-d and 6f-h). Both 5-HT-lir and FaRP-lir neurites had Syn-lir puncta along their length at non-terminal sites, which suggests the formation of *en-passant* synapses at multiple points along a single neurite [50] (Figures 6b-d and f-h). 5-HT-lir neurites, however, were more likely to have Syn-lir puncta at the end of a strand of 5-HT-lir, which resembled terminal boutons (Figures 6b-d). Syn-lir puncta were occasionally observed in close proximity to FaRP-lir blood vessels in the light organ lobe (Figure 6h, open arrowheads), which may point to light organ vasculature as an effector tissue targeted by LP neurites. Interestingly, Syn-lir puncta near blood vessels were not accompanied by FaRP-lir neurites, suggesting that these synapses belong to cells that contain either 5-HT or a yet unidentified neuroactive substance.

### 2.6 The anterior appendage is sparsely innervated from the lobe plexus

Immunostaining for 5-HT and Syn also revealed a subset of neurites originating from the LP that invade the anterior appendages (Figures 7a and d). Out of 10 anterior appendages sampled, five contained multiple 5-HT-lir neurites that terminated within the basal half of the appendage (basal appendage neurites) and a single 5-HT-lir neurite that projected notably farther (apical appendage neurite), terminating in the apical half of the anterior appendage (Figure 7a, open and closed arrowheads respectively). Of the remaining five sampled appendages, four contained only basal appendage neurites (Additional file 1: Figure S2a), and one had no detectable appendage neurites (Additional file 1: Figure S2b). 5-HT-lir in appendage neurites, like in LP neurites, was not contiguous, making the exact number of basal appendage neurites difficult to quantify using the current data. When present, the apical appendage neurite appeared to tangentially contact unlabeled cells in the appendage, but we did not observe perinuclear 5-HT-lir that would identify them as the neuron cell body from which the apical appendage neurite originates (Figure 7d). Rather, this neurite exhibits Syn-lir puncta where it tangentially contacts unlabeled cells (Figure 7d, arrowheads), again suggesting the formation of *en-passant* synapses with yet unidentified target cells in the appendage.

**FIGURE 7.**
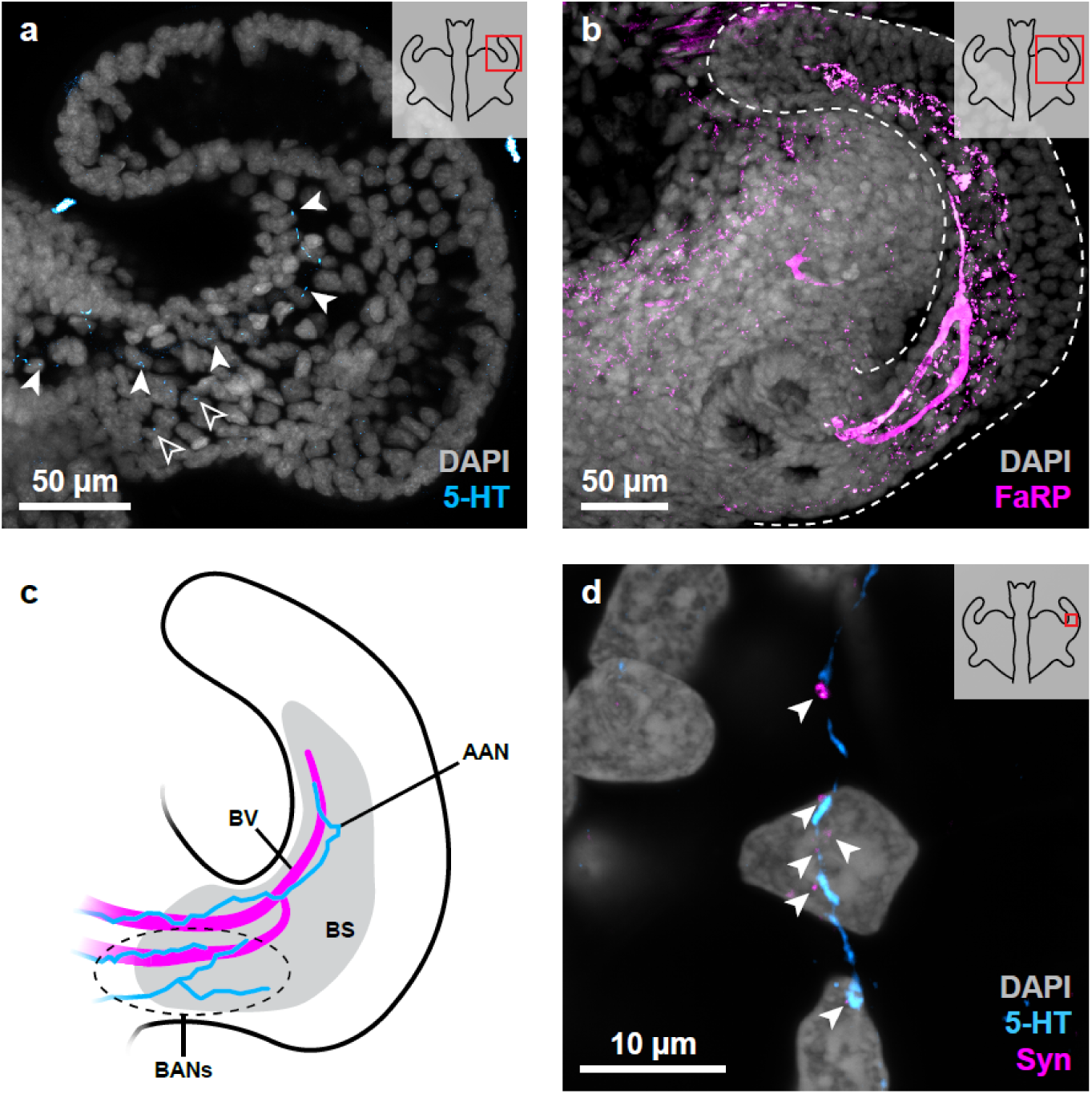
Innervation of the anterior appendage. (**a**) The anterior appendage is innervated by multiple serotonin-like immunoreactive (5-HT-lir, cyan) neurites near its base (open arrowheads) and a single 5-HT-lir neurite that projects toward the tip of the anterior appendage. (**b**) Blood vessels (BVs) and the blood sinus (BS) are revealed by FMRFamide-related peptide (FaRP, magenta) immunostaining. Dotted outline indicates anterior appendage. (**c**) Schematic diagram depicting the spatial overlap between the BVs, basal appendage neurites (BANs), and apical appendage neurite (AAN) within the BS. (**d**) The AAN appears to tangentially contact cells in the anterior appendage and displays a pattern of synapsin-1 (Syn)-lir (magenta, closed arrowheads) resembling *en-passant* synapses. The region of the light organ depicted in each panel is indicated by a red square in the inset schematic. Nuclei are stained with DAPI (grey). 5-HT, serotonin; AAN, apical appendage neurite; BANs, basal appendage neurites; BS, blood sinus; BV, blood vessel; FaRP, FMRFamide-related peptide; synapsin-1, Syn.

FaRP-lir revealed two converging blood vessels that also invade the anterior appendage (Figure 7b). The exact position of the blood vessels varied slightly between individuals, but they consistently enter the appendage from the lobe near the ducts and closely follow the internal edge of the appendage. The blood vessels converge into a Y-shaped junction in the basal half of the appendage and terminate in the blood sinus. Interestingly, the apical appendage neurite and blood vessels travel similar paths through the appendage (Figure 7c), though this could not be shown directly due to the shared host species of the primary antibodies used to visualize them. The apparent spatial overlap between innervation and vasculature in the anterior appendage may allude to a functional association between these two structures, but this remains to be determined.

## 3 Discussion

The results of this study serve to characterize the physical and neurochemical landscape of the LONS in *E. scolopes* hatchlings for the first time. We show that the LONS is a structurally elaborate segment of the PNS that displays prominent 5-HT-like and FaRP-like immunoreactivity, morphologically diverse neurons, and evidence of chemical synapses.

### 3.1 Neurite organization in the LONS

The LONS contains a strikingly complex arrangement of neurite bundles that distribute fine innervation throughout the light organ. The VMNs and VC are the largest neurite bundles in the LONS, and their organization allows for multiple possible routes for an individual neurite to enter, exit, or traverse the geography of the light organ. Both VMNs appear to enter the light organ anterodorsally and are connected by the VC near their respective hila (Figures 2 and 8a-b). Some of the neurites that compose each VMN may use the VC to innervate the contralateral half of the light organ, while the remaining neurites innervate the ipsilateral half (Figure 8c). Alternatively, each VMN may exclusively innervate the light organ contralaterally (Figure 8d). The bilateral segregation of neurites innervating the light organ raises the possibility that the LONS may originate from multiple discrete populations of neurons.

**FIGURE 8.**
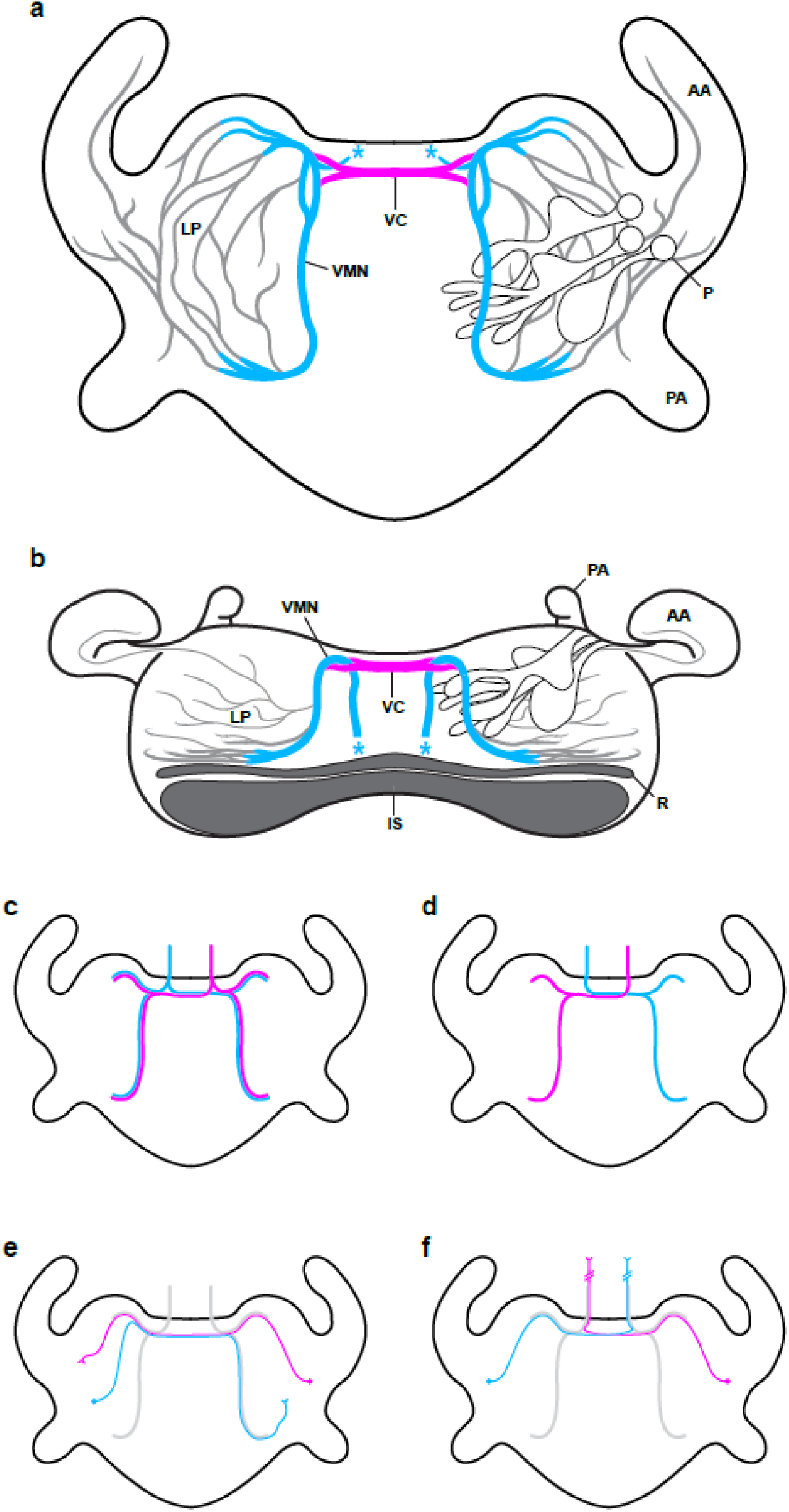
Summary of light organ-associated nervous system organization. (**a-b**) Illustrations of major light organ-associated nervous system (LONS) features from the ventral (**a**) and anterior (**b**) aspects. The ventral commissure (VC, magenta), ventromedial nerves (VMNs, cyan), and lobe plexuses (LPs, grey) are depicted in relation to key internal light organ structures. Asterisks represent the location of putative hila where the VMNs invade the light organ. (**c-f**) Schematic diagrams of possible projection pathways through the LONS. Colors indicate the side of the nervous system neurites originate from (cyan, left; magenta, right). (**c**) VMN neurites may innervate the light organ ipsilaterally and contralaterally. (**d**) VMN neurites may exclusively innervate the light organ contralaterally. (**e**) LP neurons may travel through the VC to access a target in the contralateral lobe. (**f**) LP neurons may cross the midline before exiting the light organ to innervate a distant contralateral target. AA, anterior appendage; IS, ink sac; LP, lobe plexus; P, pore; PA, posterior appendage; R, reflector; VC, ventral commissure; VMN, ventromedial nerve.

The presence of two separate anterodorsal hila also supports the majority of LONS innervation being extrinsic, which is further evidenced by the abundance of neurites and comparatively small number of NeuN-lir cell bodies (Figure 3b) and Golgi-Cox^+^ neurons (Figure 4) in the light organ. The complete projection paths and intended targets of LONS-originating neurons, however, remains unclear. One possibility is that these cells send axons through the VC to locally connect neurons in each half of the LONS (Figure 8e), perhaps to generate a rapid coordinated response to the detection of *V. fischeri*, or they may exit the light organ via the VMNs (Figure 8f).

### 3.2 The LP as a site of PNS refinement

The LP, representing the bulk of light organ innervation, is a dense meshwork of apparently disorganized innervation that spreads throughout the light organ lobe and exhibits widespread Syn-lir (Figures 2 and 6). PNS structures such as neuromuscular junctions and the myenteric plexus contain an excess of synaptic connections that are either eliminated or stabilized on the basis of their level of activation during a critical developmental window [51, 52]. Combined with altered nervous system-associated gene expression in the colonized juvenile light organ [28], these observations identify the LP as a candidate site for synaptic pruning and process refinement during post-embryonic development.

A growing body of literature indicates that the microbiome plays diverse roles in the development, maturation, and normal functioning of the ENS. Gut microbiota stimulate neuronal 5-HT production, which leads to the maturation of enteric neurons via 5-HT_4_R activation [14]. Conversely, mice without gut microbiomes exhibit gut motility defects attributed to abnormal post-embryonic ENS development [15]. Considering this, future research should address the possibility that LP refinement, if observed, may be affected by the presence of *V. fischeri*.

### 3.3 Innervation of ephemeral light organ structures

Notably, we identified a small number of prominent 5-HT-lir neurites that innervate the anterior appendage (Figure 7a). The apical and basal appendage neurites are of particular interest due to the fate of the structures they innervate. Approximately 4 days after being colonized by *V. fischeri*, the ciliated epithelium undergoes apoptosis, resulting in the complete regression of the appendages [24, 53]. Thus, neurites in the appendage must also be eliminated, but the mechanism by which this occurs remains unknown.

Extraneous axonal projections are frequently subject to local degeneration in order to refine the connectivity of neural circuits [54]. Nevertheless, constructing and decommissioning neurites is resource intensive, and the energetic cost of transmitting action potentials along axons scales linearly with distance [55], incentivizing efficiency in initial pathfinding. Further, the energy required to maintain resting membrane potential increases with membrane surface area and therefore neurite length [56]. On this basis, we infer that the apical and basal appendage neurites may have an important role in either the functioning of the appendages or the process of their decommissioning. These neurites also displayed Syn-lir in a pattern that is suggestive of *en-passant* synapse formation with target cells throughout the appendage [50]. Further research should aim to elucidate how colonization by *V. fischeri* and subsequent light organ morphogenesis impacts synapse maintenance and elimination, as well as neurite retraction.

### 3.4 Serotonin and FMRFamide-related peptides in the LONS

We observed abundant and widespread 5-HT-lir and FaRP-lir throughout the LONS using immunocytochemistry (Figure 5). Each neuroactive substance was localized to distinct regions of the light organ, indicating that LONS neurites are neurochemically heterogeneous. Neurites in the VMNs, VC, and LP were strongly 5-HT-lir, but we did not observe substantial FaRP-lir in either the VMNs or the VC. Though we observed both 5-HT-lir and FaRP-lir neurites in the LP (Figures 5a and c) and appendages (Figure 7), we could not confirm whether these neuroactive substances were both present in a single cell due to the shared hosts of the antibodies used in this study. 5-HT and FMRFamide have been shown to occupy associated neuronal populations in other invertebrates [57, 58]. This suggests that the LONS may contain two distinct populations of neurons expressing 5-HT and FaRPs respectively, though the functional roles of each neuroactive substance remain unexplored.

FaRP-lir was localized within LONS neurites, as well as vascular structures in the light organ, including blood vessels in the lobes and appendages as well as the appendage blood sinuses (Figures 5c-d and 7b). The tetrapeptide FMRFamide was first described as a cardioexcitatory peptide in *M. nimbosa* [46]. Since then, FaRPs have been identified in all major invertebrate phyla and are implicated in numerous physiological processes, including but not limited to neuronal activity, circulatory regulation, and neuroimmune interactions [47]. Our observations of FaRP-lir in both neuronal and vascular tissues point to a multifaceted role for FaRPs that may involve neuromodulation and regulation of blood flow within the light organ, as well as signaling between the LONS and *V. fischeri*.

5-HT is a widely utilized bioamine in ENS tissues and is critical for governing gut motility via peristaltic contraction in both vertebrates [59, 60] and invertebrates [61, 62]. Adult *E. scolopes* expel up to 90% of their resident symbiont population on a diel rhythm by contracting the light organ [63, 64], but the systems that control symbiont venting remain unknown. Given the light organ’s close relation to the gut and abundance of 5-HT-lir neurites, we propose contraction controlled by serotonergic LONS innervation as a candidate mechanism for future investigation.

It should also be noted that our results pertaining to the neuroactive substances in the LONS are not exhaustive. The existence of other classical neuroactive substances (glutamate, γ-aminobutyric acid, acetylcholine, and dopamine) as well as those common in invertebrates (octopamine, tachykinin-related peptides, and other RFamides) have not been scrutinized in the LONS and should be the subject of future investigations.

### 3.5 Putative intrinsic LONS neurons

Using an antibody against NeuN, we were able to locate putative neuron cell bodies within the LONS (Figure 3a). NeuN-lir cells were most frequently observed near the ducts, antechambers, and deep crypts (Figures 3c-e). Our novel Golgi-Cox staining protocol tailored for *E. scolopes* hatchlings supported our NeuN findings and additionally revealed an assortment of cells in the light organ whose morphology strongly resembled unipolar, bipolar, and multipolar neurons [42] (Figure 4). However, it cannot be ruled out that a subset of Golgi-Cox^+^ cells are instead peripheral glia or another yet undetermined non-neuronal PNS cell type [65]. Further investigation of LONS cell type diversity is needed to address this possibility. It should also be noted that the Golgi-Cox method is generally agreed to reveal fewer than 5% of cells in a given sample [40, 66, 67]. Consequently, the results of our Golgi-Cox experiments in isolation cannot be used to draw firm conclusions about the neuronal population makeup of the LONS. Rather, they aptly demonstrate the morphological diversity of LONS cells and the utility of our novel staining protocol for further interrogation of the LONS. In any case, these putative neurons are uniquely positioned within the light organ to receive signals from the epithelial cells that compose the ISS and directly interface with *V. fischeri*. In response to bacterial products, mammalian enterochromaffin cells produce 5-HT, which is released to locally activate 5-HT receptors on enteric neurons to induce a range of physiological changes [9, 59, 60], supporting a potential role for LONS neurons in the detection of symbionts within the light organ.

## 4 Conclusion

Here, we present a detailed description of the organization and neurochemical landscape of the LONS of hatchling *E. scolopes* that lays the groundwork for future research investigating LONS function and physiology, as well as how a single bacterial symbiont can drive post-embryonic PNS development in its host. Our data suggest that the LONS is positioned to play a key role in the initiation, establishment, and maintenance of the squid-*Vibrio* symbiosis, as well as the mediation of canonical light organ morphogenesis [17, 18, 24]. From the neurochemical and morphological diversity of LONS neurons, we infer underlying transcriptional diversity which will prove valuable in developing techniques for targeting and manipulating these cells. Additionally, the apical appendage neurite provides a uniquely powerful opportunity to study the mechanisms by which bacterial MAMPs impact the fate of a single, easily identifiable neuronal process. Overall, our initial observations of the *E. scolopes* LONS provide a new foundation from which to interrogate the dynamic interactions between beneficial bacterial symbionts and host nervous systems.

## 5 Methods

### 5.1 Animals

Adult Hawaiian bobtail squid *(Euprymna scolopes)* were collected by dipnet from Maunalua Bay, O’ahu, Hawai’i and transported to East Lansing, Michigan. The adult animals were housed in individual compartments of a 950 L recirculating aquarium with artificial seawater (ASW) made using made using Instant Ocean (Spectrum Brands, Inc.). Females were provided sections of PVC pipe to deposit fertilized egg clutches, and soon after being deposited, clutches were removed from the aquarium compartments and placed in a 285 L recirculating aquarium with ASW that does not contain *Vibrio fischeri*. Hatchling squid were collected from the aquarium and anaesthetized in 2% ethanol in filter-sterilized ASW within 3 h of hatching. Hatchlings were then fixed with 4% paraformaldehyde in marine phosphate-buffered saline (mPBS; 100mM sodium phosphate, 0.9 M sodium chloride, pH 7.4) for 24 h at 4°C and stored in mPBS at 4°C. All experiments were conducted in compliance with Michigan State University Institutional Animal Care and Use Committee standards.

### 5.2 Immunocytochemistry

An anti-5-HT polyclonal antibody (1:1000; ImmunoStar, 20080; RRID:AB_572263) was used to broadly visualize LONS architecture (*n* = 10). Putative neuron cell bodies in light organs (*n* = 10) were labeled using an anti-NeuN polyclonal antibody (1:100; Millipore, ABN78C3; RRID:AB_11204707). We used a rabbit anti-FMRFamide polyclonal antibody (1:1000; Millipore, AB15348; RRID:AB_805291) to label FMRFamide and to a lesser degree other FaRPs, a family that includes numerous invertebrate neuropeptides [47, 68]. To label presynaptic neuron terminals in combination with 5-HT (*n* = 10) and FaRPs (*n* = 10), we used mouse anti-SYNORF1 (1:20; DSHB 3C11, deposited by Buchner, E.; RRID:AB_528479, Developmental Studies Hybridoma Bank), which recognizes a 10-aa epitope in the conserved C-terminal domain of synapsin-1 (Syn), a synaptic vesicle binding protein. Clone 3C11 has been used to visualize Syn in the pelagic squid *Abraliopsis falco* [69], and a similar pattern of staining was observed in the LONS. For all antibodies, specific binding in the LONS was confirmed by omitting primary antibodies, which showed no labeling of LONS elements. This ensured that immunofluorescent labeling was not the result of non-specific binding of secondary antibodies or autofluorescence.

For all immunocytochemistry, light organs were dissected from fixed hatchling animals under a Leica S9i stereomicroscope. Ink sacs were punctured during dissection, and light organs were washed in mPBS to remove any remaining ink, then washed in mPBS-Triton X-100 (mPBST, 1%). Light organs were blocked for 24h using bovine serum albumin (BSA, 5%; Invitrogen, 15561020) and normal goat serum (1%; Invitrogen, 31837) in mPBST, followed by incubation in primary antibody diluted in blocking solution for 7 days. After primary antibody incubation, samples were washed in mPBST and blocked for 24 h. Light organs were then incubated in secondary antibodies diluted in blocking solution for 3 days. Secondary antibodies used in this study were goat anti-rabbit AlexaFluor 488 (1:750; Invitrogen, A11008; RRID:AB_143165), goat anti-rabbit AlexaFluor 555 (1:750; Invitrogen, A21428; RRID:AB_2535849), goat anti-mouse AlexaFluor 488 (1:750; Invitrogen, A11001; RRID:AB_2534069), goat anti-mouse AlexaFluor 568 (1:750; Invitrogen, A11004; RRID:AB_2534072), and goat anti-rat AlexaFluor 647 (1:750; Invitrogen, A21247; RRID:AB_141778). Samples were then washed sequentially in mPBST and mPBS, followed by an incubation in DAPI dilactate (10ug/mL in mPBS; Invitrogen, D3571) for 24 h. Following final mPBS washes, light organs were mounted in VectaShield Antifade Mounting Medium (Vector Labs, H-1000-10). Mounted samples were stored at 4°C and protected from light until imaging.

### 5.3 Golgi-Cox Staining

A novel Golgi-Cox staining protocol was optimized for use in juvenile *E. scolopes* light organs based on findings from Ranjan and Mallick [39], Zaqout and Kaindl [40], and Vints et al. [41]. Golgi-Cox impregnation solution was prepared in advance of staining by combining 50 mL of 5% potassium dichromate, 50 mL of 5% mercury chloride, 40 mL of 5% potassium chromate, and 100 mL DI H_2_O (all solutions w/v in DI H_2_O). The solution was protected from light and maintained at room temperature for 48h to allow for precipitate formation. For impregnation, PFA-fixed hatchling animals were placed in 5 mL Golgi-Cox solution in scintillation vials for 48 h. During impregnation, scintillation vials were protected from light and kept in a water bath to heat the solution to 37°C. After 24 h, samples were transferred into a new scintillation vial containing fresh Golgi-Cox solution for the remainder of the 48h impregnation period. To develop the stain, samples were removed from Golgi-Cox solution and serially washed in the following: DI H_2_O twice for 5 min each; 28% ammonium hydroxide for 8 min; DI H_2_O twice for 5 min each; Kodafix (B&H Photo, KOFXRPDR) for 30 min; DI H_2_O twice for 5 min each; 50% EtOH twice for 5 min each; 70% EtOH twice for 5 min each; 95% EtOH twice for 5 min each; 100% EtOH twice for 5 min each; xylene twice for 5 min each. For all developing steps, samples were protected from light and gently agitated. After developing, light organs were dissected out and mounted in DPX (Sigma Aldrich, 06522), stored at 4°C, and protected from light until imaging. Staining was deemed successful by positive staining of at least one cell with neuron-like morphology, and all successfully stained light organs were considered in the analysis (*n* = 15).

### 5.4 Imaging and Image Analysis

Lightfield images of hatchling animals and whole light organs were captured using a Leica THUNDER Imager Model Organism with a Leica K5 camera. Whole light organ immunofluorescent stains were imaged in the MSU Advanced Microscopy Core (RRID:SCR_027702) using a Leica Stellaris 5 confocal microscope (Leica Microsystems, RRID:SCR_024663) running Leica Application Suite X (Leica Microsystems, RRID:SCR_013673). Optical sections were captured and digitally combined to form maximum intensity projections for analysis. Images of Golgi-Cox-stained light organs were taken using a Keyence VHX-6000 digital microscope (Keyence Corporation). All fluorescence micrographs collected were given false color and adjusted for brightness and contrast using FIJI (version 2.16.0/1.54p, National Institutes of Health, RRID:SCR_002285).

### 5.5 NeuN-lir Cell Kernel Density Estimate

To construct a distribution map of NeuN-lir cell bodies, Z-stacks (step size = 2 μm) of whole light organs (*n* = 10) were captured at 200x magnification such that the entire organ is in frame. In FIJI, Z-stacks were rotated so the anterior-posterior axis was vertical, using the top edge of the organ for alignment. Choosing the confocal section containing the widest plane of the light organ, Z-stacks were then cropped in the X and Y directions such that each edge of the image was tangent to the edge of the organ, making the image dimensions equal to the maximum width and height of the light organ. NeuN-lir cells were then plotted using the Multipoint tool, placing the marker at the center of each cell’s nucleus. The X and Y coordinates as well as the confocal section number (Z) of each cell observation were recorded as a proportion of the total width, height, and depth of the light organ to account for variation in light organ size between individuals. Normalized light organ coordinates were then used to generate two-dimensional kernel density estimates (2D-KDEs) using the MASS package (v. 7.3-65, RRID:SCR_019125) in R (v. 4.5.0, RRID:SCR_001905). 2D-KDEs were computed using axis-aligned bivariate Gaussian kernels with a bandwidth (*h*) of 0.1, applied in both directions for all plots. Bandwidth was manually selected to optimize bias and variance while considering the size of a NeuN-lir nucleus relative to the whole light organ.

NeuN-lir cell observations in the appendages were included in total cell counts but were omitted from the 2D-KDEs due to the variable position of the appendages during imaging. Total counts were plotted using GraphPad Prism (v. 10.6.1 for Windows, GraphPad Software, Boston, Massachusetts USA; RRID:SCR_002798).

## Declarations

### Ethics approval and consent to participate

All experiments were conducted in compliance with Michigan State University Institutional Animal Care and Use Committee standards (08/24-005-99).

### Consent for publication

Not applicable

### Availability of data and materials

The datasets used and analyzed during the current study are available from the corresponding author on reasonable request.

### Authors’ Contributions

ABW designed and performed all experiments, analyzed the resulting data, and drafted the manuscript. EVXW optimized experimental protocols and contributed to experimental design. EACH-H drafted and revised the manuscript, contributed to experimental design, and assisted with data interpretation. All authors contributed to the final writing of the manuscript.

## Acknowledgements

The animals used in our experiments were collected from Maunalua Bay, which is located in the ahupuaʻa (land division) of Waimānalo, in the Pae ʻĀina o Hawaiʻi (Hawaiian archipelago). Hawaiʻi is an Indigenous space without which our research would not be possible. We express our gratitude to the Kānaka Maoli (Native Hawaiians) for their hospitality and systems of knowledge, which have sustained Hawaiʻi and allowed generations of scientists to learn from this ʻāina (land) and its people. All imaging was performed at the Michigan State University Center for Advanced Microscopy Core Facility (RRID:SCR_027702). The monoclonal antibody developed by E. Buchner (Universitaetsklinikim Wuerzburg, Institute of Clinical Neurobiology) was obtained from the Developmental Studies Hybridoma Bank, created by the NICHD of the NIH and maintained at The University of Iowa, Department of Biology, Iowa City, IA 52242. Finally, we thank Dr. Julia Ganz for her discussion of the manuscript.

## Authors’ information

Not applicable

## Footnotes

Not applicable

## Supplementary Information

Additional file 1: Supplemental Figures S1 and S2 (LONS_Supplementary-Information.pdf)

**Figure S1** | Scatter plots showing NeuN-like immunoreactive (-lir) cells in single light organs. (**a**) Normalized coordinate plane for ventral view (frontal plane) of the light organ. Red box bounding the light organ diagram corresponds to the extreme values of the coordinate planes depicted in (**b-k**). (**b-k**) Scatter plots showing the location of NeuN-lir cells in the frontal plane of 10 light organs. Observations from these individuals were aggregated to perform the 2-dimensional kernel density estimates in Figures 3d-e.

**Figure S2** | Alternate patterns of anterior appendage innervation. (**a**) Anterior appendage with serotonin-like immunoreactive (5-HT-lir, cyan) basal appendage neurites (BANs) but lacking a 5-HT-lir apical appendage neurite (AAN). (**b**) Anterior appendage lacking an AAN and BANs. Nuclei are stained with DAPI (grey). 5-HT, serotonin; AA, anterior appendage.

