## Supplementary material for "The Neuroanatomy of the Hawaiian Bobtail Squid Juvenile Bacterial Light Organ": Suppl

### Supplementary Information

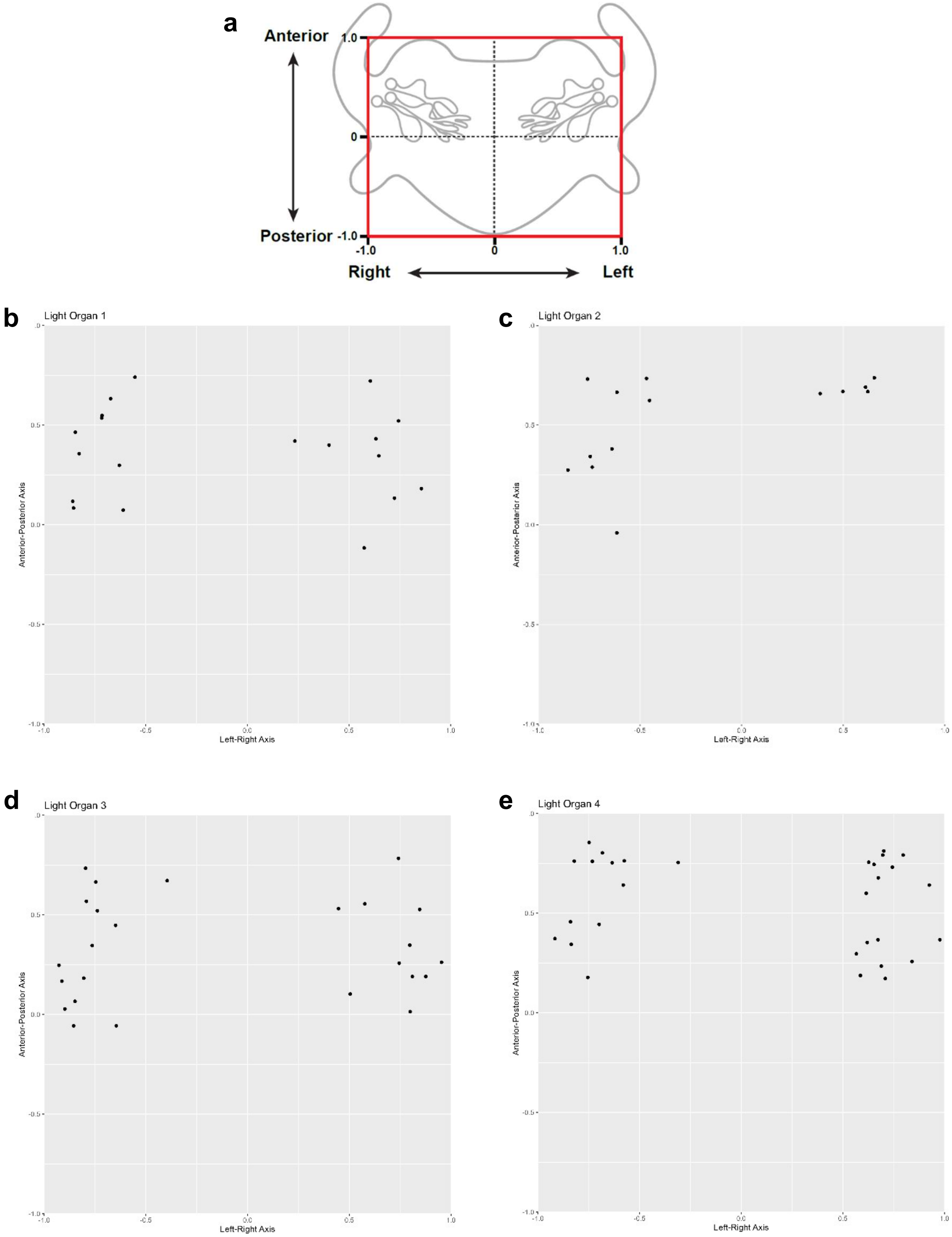

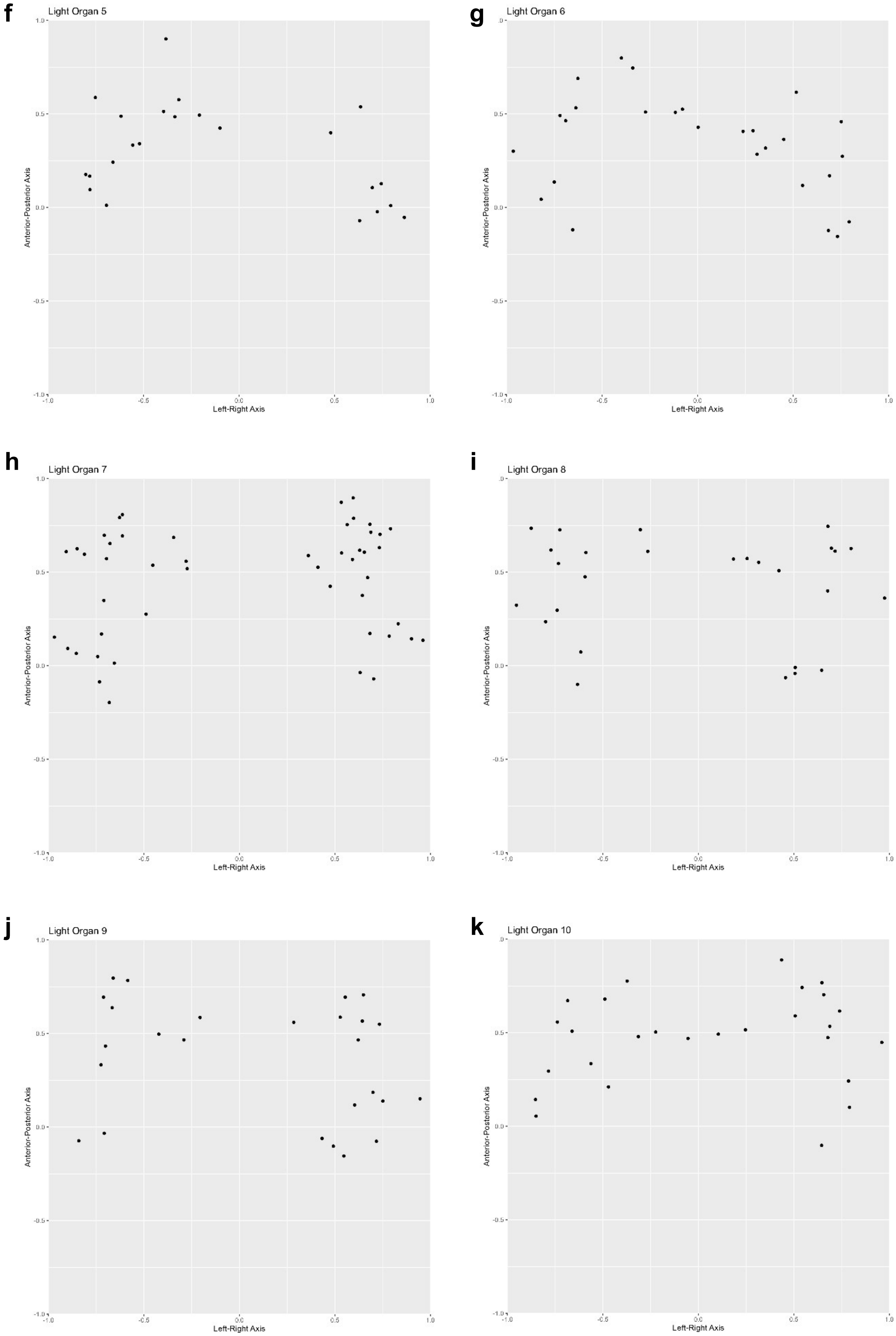
f
Light Organ 5
g
Light Organ 6
h
Light Organ 7
i
Light Organ 8
j
Light Organ 9
k
Light Organ 10
Anterior-Posterior Axis
Left-Right Axis
1.0
0.5
0.0
-0.5
-1.0

**Figure S1 |** Scatter plots showing NeuN-like immunoreactive (-lir) cells in single light organs. (**a**) Normalized coordinate plane for ventral view (frontal plane) of the light organ. Red box bounding the light organ diagram corresponds to the extreme values of the coordinate planes depicted in (**b-k**). (**b-k**) Scatter plots showing the location of NeuN-lir cells in the frontal plane of 10 light organs. Observations from these individuals were aggregated to perform the 2-dimensional kernel density estimates in Figures 3d-e.

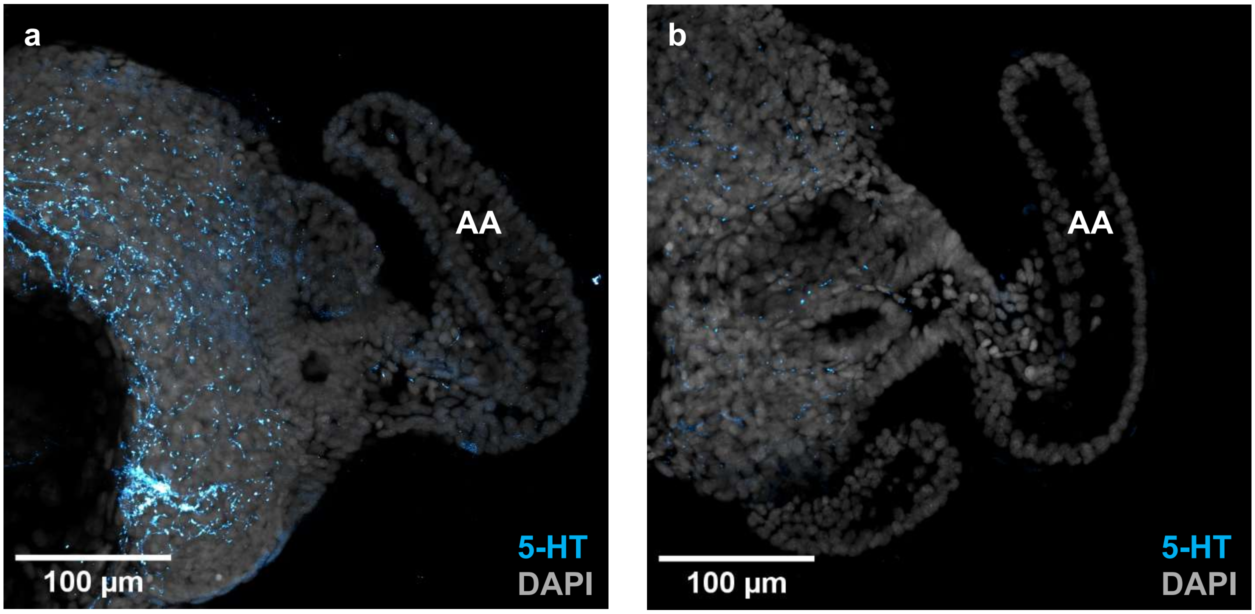


**Figure S2 |** Alternate patterns of anterior appendage innervation. (**a**) Anterior appendage with serotonin-like immunoreactive (5-HT-lir, cyan) basal appendage neurites (BANs) but lacking a 5-HT-lir apical appendage neurite (AAN). (**b**) Anterior appendage lacking an AAN and BANs. Nuclei are stained with DAPI (grey). 5-HT, serotonin; AA, anterior appendage.
